# Targeting the CAPON–NOS Axis: A Computational Strategy for Small Molecule Modulator Discovery

**DOI:** 10.1101/2025.08.05.668619

**Authors:** Hossam Nada, Gerhard Wolber, Moustafa T. Gabr

## Abstract

The carboxy-terminal PDZ ligand of neuronal nitric oxide synthase (CAPON) serves as a critical regulatory protein controlling nitric oxide (NO) signaling across multiple physiological and pathological processes which encompass neurological, cardiac and metabolic functions. These diverse physiological roles of CAPON marks it as a key therapeutic target for conditions associated with its dysregulation. Despite this therapeutic potential there are no specific CAPON or nNOS/CAPON modulators which have been developed to date, highlighting a significant gap in targeted drug discovery. Herein, we report the first strategy specifically focused on disrupting the nNOS/CAPON protein-protein interface. Through screening of chemical libraries composed of 4.6 million compounds and eight molecular dynamics simulations, two potential hit compounds were identified.

Beyond identifying these promising hits, our approach introduces two novel computational tools: a freely available Python-based toolset for NMR structural analysis and visualization and a second toolkit for accelerated ligand preparation. These tools significantly accelerate data preparation timelines while reducing computational costs, providing the research community with accessible resources for structure-based drug discovery efforts. Together, these tools represent a substantial contribution to the computational chemistry toolkit, enabling researchers to conduct high-throughput virtual screening campaigns more efficiently and with greater reproducibility. This work represents a foundational step toward developing targeted therapies for CAPON-mediated disorders and provides a scalable computational framework for future protein-protein interaction drug discovery efforts.

**Highlights:**

- A novel structure-based strategy developed to target the CAPON/nNOS protein–protein interaction.
- A Python-based toolset for protein conformation analysis, identification, visualization and separation.
- Python pipeline enables efficient ligand preparation for ultra-large chemical libraries.
- Virtual screening identified 6 promising small-molecule candidates.
- A total of 8*100ns molecular dynamics (MD) simulations performed using DESMOND.
- MM/GBSA and contact-time analysis were conducted to assess binding stability and affinity.

## 1. Introduction

The carboxy-terminal PDZ ligand of neuronal nitric oxide synthase (CAPON) is a key regulatory protein which is responsible for the modulation of nitric oxide (NO) signaling in various physiological and pathological processes.^1, 2^ Nitric oxide synthase (NOS) exists in three distinct isoforms: neuronal-type (nNOS), inducible-type (iNOS), and endothelial-type (eNOS)^3^. Each of the NOS isoforms serve distinct specialized functions across different tissues including the cerebellum, skeletal muscles, kidneys, blood vessels, and immune cells^4–6^. Among the nine known proteins that interact with nNOS, CAPON is a crucial regulator of nNOS activity through its unique ability to compete with postsynaptic density protein 95 (PSD-95) for binding to the nNOS PDZ domain^7^. This competitive interaction fundamentally alters NO production and downstream signaling cascades, positioning CAPON as a master regulator of nitric oxide homeostasis.

The CAPON signaling pathway (Figure 1) operates through a complex molecular switching mechanism that directly impacts neuronal function and cellular metabolism. Under normal conditions, nNOS forms a functional complex with N-methyl-D-aspartate (NMDA) receptors and PSD-95, facilitating calcium-dependent NO production in response to glutamatergic signaling^8^. However, CAPON disrupts this canonical NMDAR-PSD95-nNOS complex by competing for the same PDZ-binding motif on nNOS, resulting in the formation of an alternative NMDAR-CAPON-nNOS complex^2, 9^. This molecular rearrangement significantly attenuates nNOS activity and reduces NO production, thereby providing neuroprotection against excitotoxicity and oxidative stress. The downstream effects of CAPON-mediated nNOS regulation extend beyond the nervous system, influencing calcium channel function in cardiac myocytes^10^ through S-nitrosylation of L-type calcium channels and modulating insulin signaling pathways^11, 12^ in pancreatic cells. This multifaceted signaling network positions CAPON as a central hub connecting neuronal activity, cardiac electrophysiology, and metabolic regulation.

**Figure 1.**
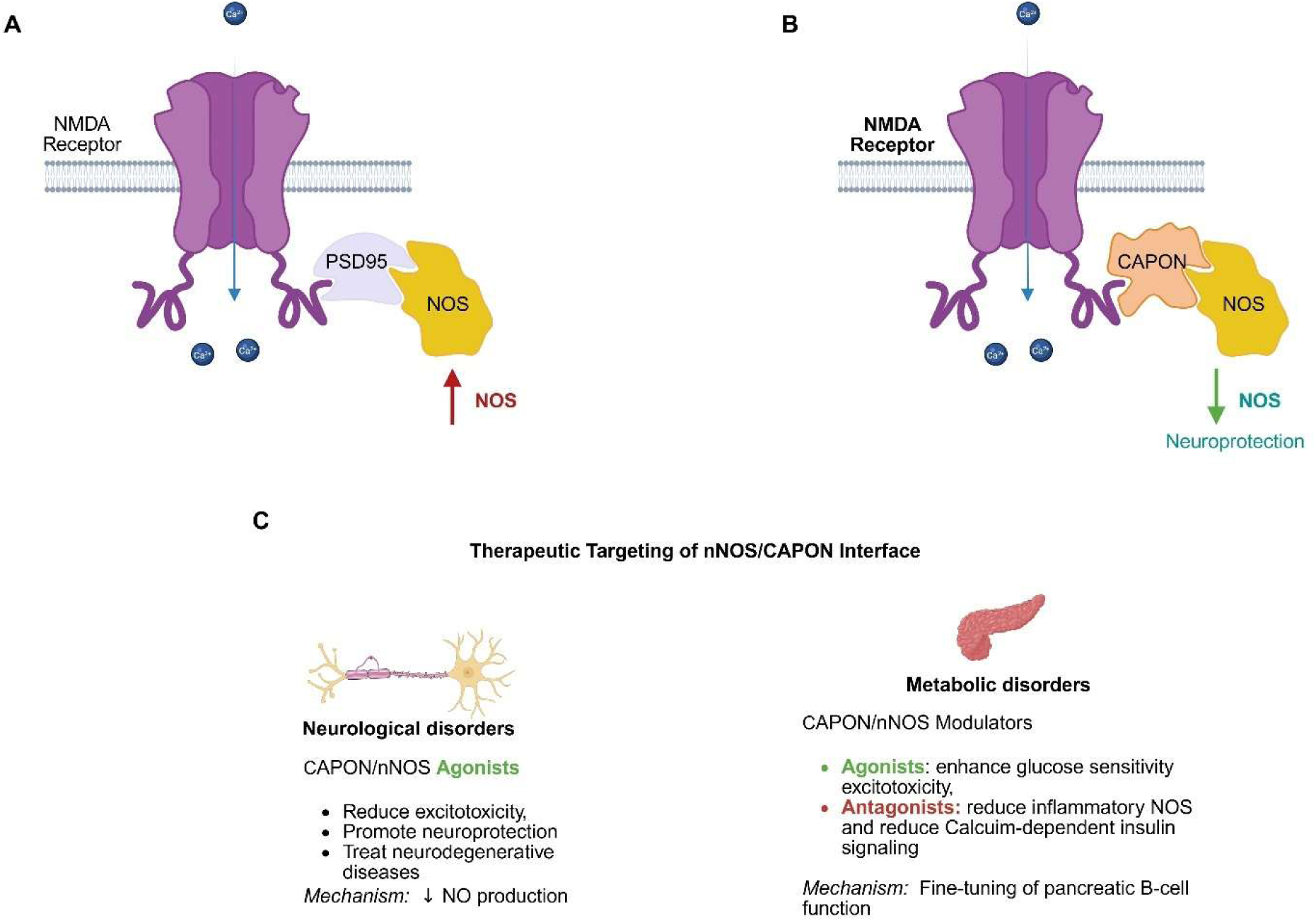
CAPON-Mediated Modulation of NMDA Receptor–nNOS Signaling and Its Therapeutic Potential. (A) Under normal physiological conditions, the activation of NMDA receptors lead to calcium influx which promotes the recruitment of PSD95 and neuronal nitric oxide synthase (nNOS), resulting in high nitric oxide (NO) production. This Ca²⁺-dependent NO signaling plays roles in neurotransmission but can also contribute to excitotoxicity and neuronal damage. (B) CAPON competitively binds to nNOS, displacing PSD95 and disrupting the NMDA receptor–nNOS signaling complex. This reduces NO production and confers neuroprotective effects by attenuating excitotoxicity. (C) Therapeutic strategies targeting the CAPON–nNOS interface include the use of agonists and modulators: In neurological disorders, CAPON/nNOS agonists reduce excitotoxicity and promote neuroprotection by decreasing NO production. In metabolic disorders, CAPON/nNOS modulation via agonists and antagonists are theorized to have different therapeutic effects. CAPON agonists will enhance glucose sensitivity and modulating excitotoxicity. Conversely, CAPON antagonists will reduce inflammatory NO production and restoring calcium-dependent insulin signaling.

The therapeutic potential of CAPON is highly context-dependent due to its complex signaling mechanism. This is due to the diverse roles of CAPON which often necessitate opposing pharmacological approaches depending on the target tissue and disease context. In neurological disorders (Figure 1C), the neuroprotective function of CAPON through nNOS modulation presents opportunities for treating neurodegenerative diseases and reducing excitotoxic neuronal damage. While in metabolic disorders, the enhancement of CAPON activity may improve pancreatic β-cell function and glucose sensitivity, while simultaneously risking the promotion of inflammatory pathways in other tissues. As such, the impact of CAPON is highly context-dependent and disease specific. Given the involvement of CAPON in multiple pathological conditions and signaling pathways, it has emerged as a compelling therapeutic target. In addition, the modulation of CAPON function offers not only a promising strategy for novel treatments but also a valuable avenue to deepen our understanding of its biological effects. Despite this potential, to the best of our knowledge no CAPON or nNOS/CAPON modulators have been developed to date which underscores the urgent and unmet need for targeted drug discovery in this space.

Structure-based discovery of CAPON modulators has been hindered by the lack of an experimentally resolved crystal structure. As such, an alternate strategy was devised toward targeting CAPON via the nNOS/CAPON interface which was made possible by the availability of an NMR detailing the extended neuronal nitric oxide synthase pdz domain complexed with an associated peptide^13^

## 2. Results and Discussion

### 2.1. NMR conformations analysis

The NMR structure of nNOS bound in complex with Asp–X–Val–COOH peptides of CAPON contains 15 distinct conformations. Analysis of the 15 nNOS conformations revealed significant structural diversity within the backbone atoms where the pairwise RMSD values ranging from 1.19 Å to 7.18 Å (mean: 3.42 ± 1.52 Å). The 8^th^ conformation (Model 8) was identified as the central conformation (Figure 2) where it exhibited the lowest mean RMSD (2.28 Å) relative to all other structures which is an indicator of its representative nature within the ensemble. K-means clustering analysis partitioned the ensemble into three distinct conformational states, with Cluster 1 containing the majority of models (73.3%), while Clusters 0 and 2 each comprised 13.3% of the structures.

**Figure 2.**
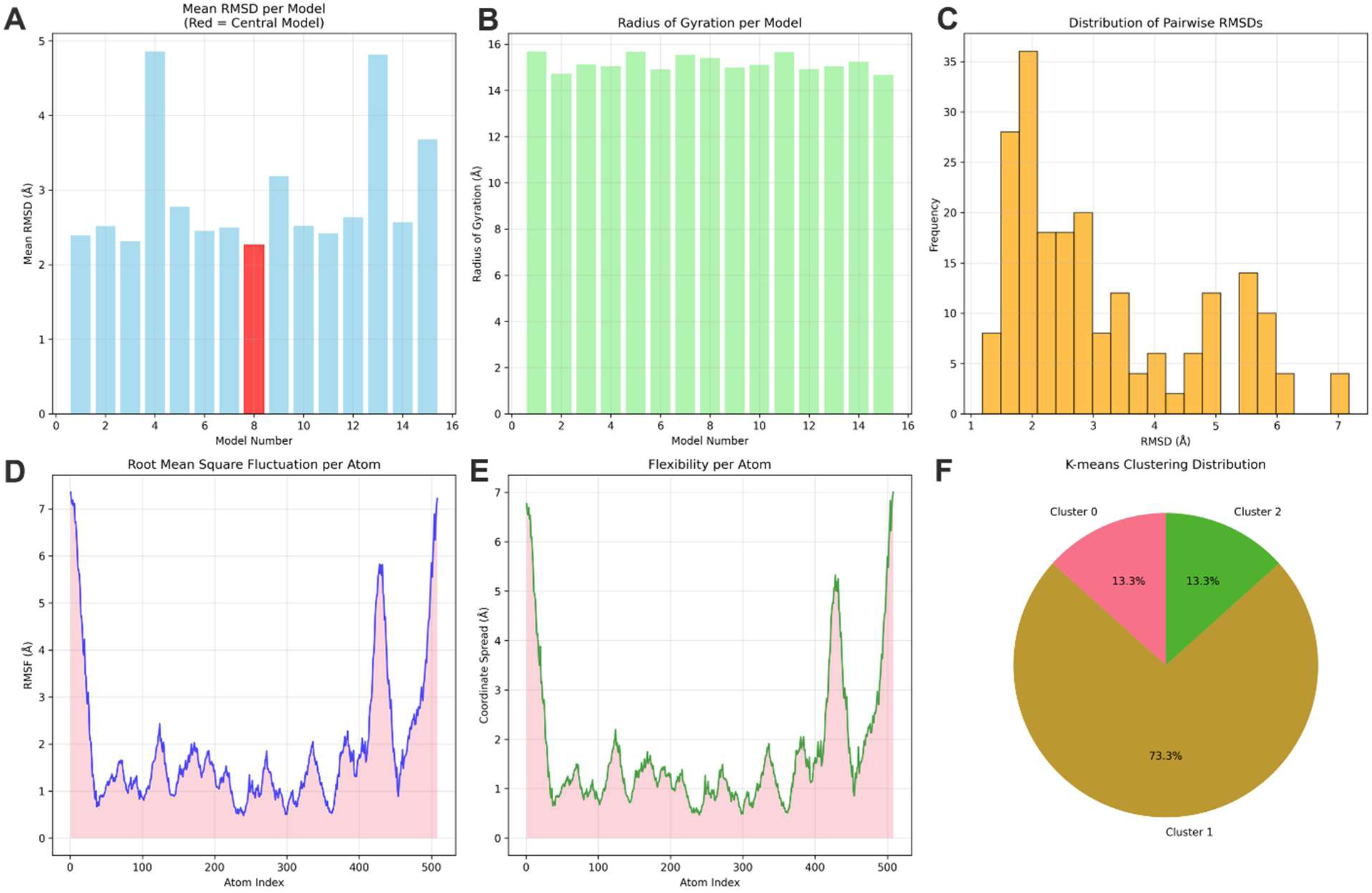
Conformational analysis summary of 15-model PDB ensemble (1B9Q, Chain A). (A) Mean RMSD per model with central model highlighted in red (Model 8). (B) Radius of gyration showing overall structural compactness. (C) Distribution of pairwise RMSD values revealing conformational diversity. (D) Root mean square fluctuation per backbone atom highlighting flexible regions. (E) Coordinate spread analysis showing per-atom flexibility. (F) K-means clustering distribution into three conformational states.

The radius of gyration values showed relatively modest variation (14.7-15.7 Å) which suggests that while local conformational changes are substantial, the overall protein compactness remains largely conserved across models. Furthermore, the narrow range of radius of gyration values indicate that the overall protein architecture remains intact and that the observed diversity represents functionally relevant motions rather than unfolding events.

Principal component analysis (Figure 3) revealed that the first three components captured 83.6% of the total conformational variance (PC1: 62.7%, PC2: 14.5%, PC3: 8.8%), with the PCA projection clearly separating models into distinct groups where outlier conformations (M4, M13, M9, M15) exhibited higher mean RMSD values (>4.0 Å) compared to the central cluster of structurally similar models. The substantial RMSD range observed further support the presence of considerable conformational flexibility exceeding typical thermal fluctuations and suggesting functionally relevant conformational transitions. The dominance of Cluster 1 (73.3% of models) suggests a primary functional state, while minority clusters may represent transient intermediate states or alternative functional conformations. Based on the results of analyzing the CAPON structure, the central conformation (model 8) was selected for carrying out the virtual screening assay.

**Figure 3.**
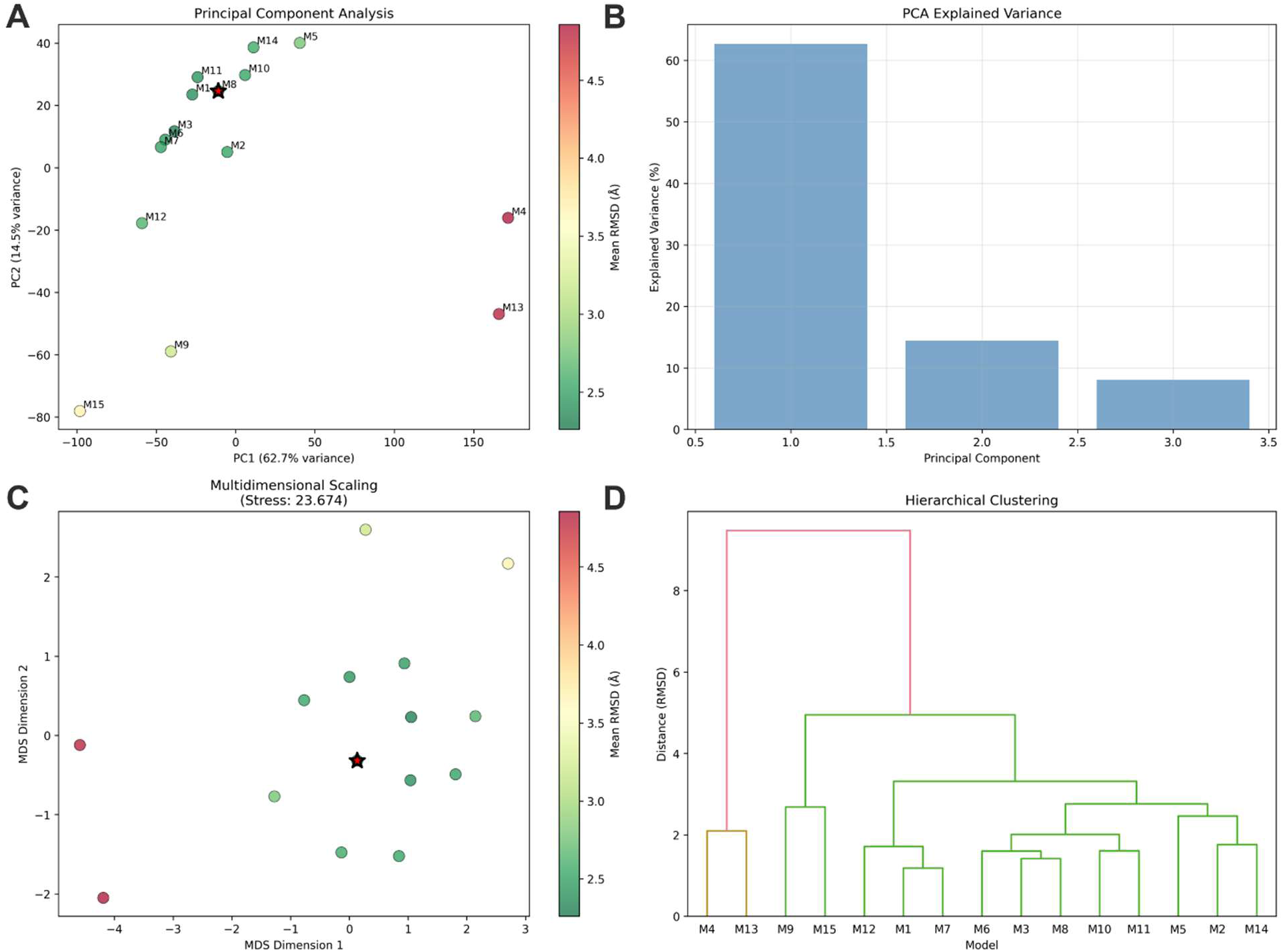
Dimensionality reduction analysis of conformational ensemble. (A) Principal component analysis projection colored by mean RMSD, with central model (black star) and model annotations. (B) PCA explained variance showing three components capture 83.6% of total variance. (C) Multidimensional scaling projection (stress: 23.674) preserving pairwise RMSD relationships. (D) Hierarchical clustering dendrogram revealing conformational families and relationships between models.

### 2.2. Library preparation

Ligand preparation for ultra-large libraries is a computationally intensive process which demands substantial computational storage and memory resources.^14, 15^ The Enamine Screening Collection (4.6 million compounds) was chosen for the screening efforts in this work due to its extensive chemical diversity, drug-like properties and its proven track record for success high-throughput virtual screening campaigns^16–18^. The library offers a broad representation of lead-like and fragment-like molecules, making it a suitable source for identifying potential hits with favorable pharmacokinetic and physicochemical characteristics. Initial ligand preparation efforts revealed that processing these libraries on our system which is equipped with an Nvidia 470 Ti GPU and 32 CPU cores would require high computational cost and time. To address this bottleneck, a streamlined ligand preparation pipeline (supplementary folder) was developed to significantly reduce computational overhead and accelerate the processing time. The ligand preparation pipeline implemented here offers a reproducible, scalable, and chemically sound approach to generating screening-ready small molecules. The process incorporates a robust suite of standardization and filtering criteria commonly accepted in early-stage drug discovery. The pipeline’s use of RDKit^19^ for SMILES parsing, descriptor calculation, and molecular embedding ensures chemical accuracy and compatibility with cheminformatics standards.

Notably, the parallelized processing of molecules using Python’s multiprocessing tools dramatically increases throughput, making the pipeline suitable for libraries containing tens of thousands of compounds. The integration of multiple structural alert filters, including PAINS^20^ and BRENK^21^, enhances the downstream screening quality by removing promiscuous and toxic scaffolds. The charge neutralization step is especially valuable for virtual screening workflows where charged ligands may show artificial binding due to force field artifacts.

Additionally, the pipeline’s modular design allows for user customization at each stage via command-line arguments (e.g., enabling 3D coordinate generation, skipping Lipinski rules, or keeping duplicates). Comprehensive logs generated at each step offer transparency, facilitate troubleshooting, and allow auditing of filtered-out compounds with rationales. Finally, by packaging the pipeline with setup scripts and standardized dependencies (as seen in setup_package.py), it can be easily installed and shared across research teams, ensuring reproducibility and ease of deployment.

### 2.3. Virtual screening results

To identify potential compounds capable of disrupting CAPON/NOS binding, the two prepared compound libraries from Enamine were subjected to a hierarchical virtual screening workflow targeting the identified CAPON/NOS binding site. The virtual screening workflow involved the Enamine Screening Collection which comprised of 4.6 million readily available compounds. The virtual screening protocol was executed in three sequential stages using the GLIDE module of Schrodinger which involved High-Throughput Virtual Screening (HTVS), Standard Precision (SP), and Extra Precision (XP) docking of the prepared libraries. In this workflow a stepwise filtration approach was employed in which only the top 10% of compounds from each stage advanced to the next stage.

The hierarchical approach successfully identified 4,750 predicted hits (see supplementary file), which exhibited a broad range of docking scores, indicating diverse binding affinities and orientations within the target binding site. To further refine the selection, the top 1% of these hits (500 compounds with the highest docking scores) underwent visual inspection. During this manual curation step, compounds were evaluated based on their spatial orientation and binding mode relative to the native CAPON-CT7 peptide (the C-terminal 7-residue peptide) at the nNOS binding site (Figure 4A-B). This comparative analysis ensured that selected compounds would mimic the natural protein-protein interaction.

**Figure 4.**
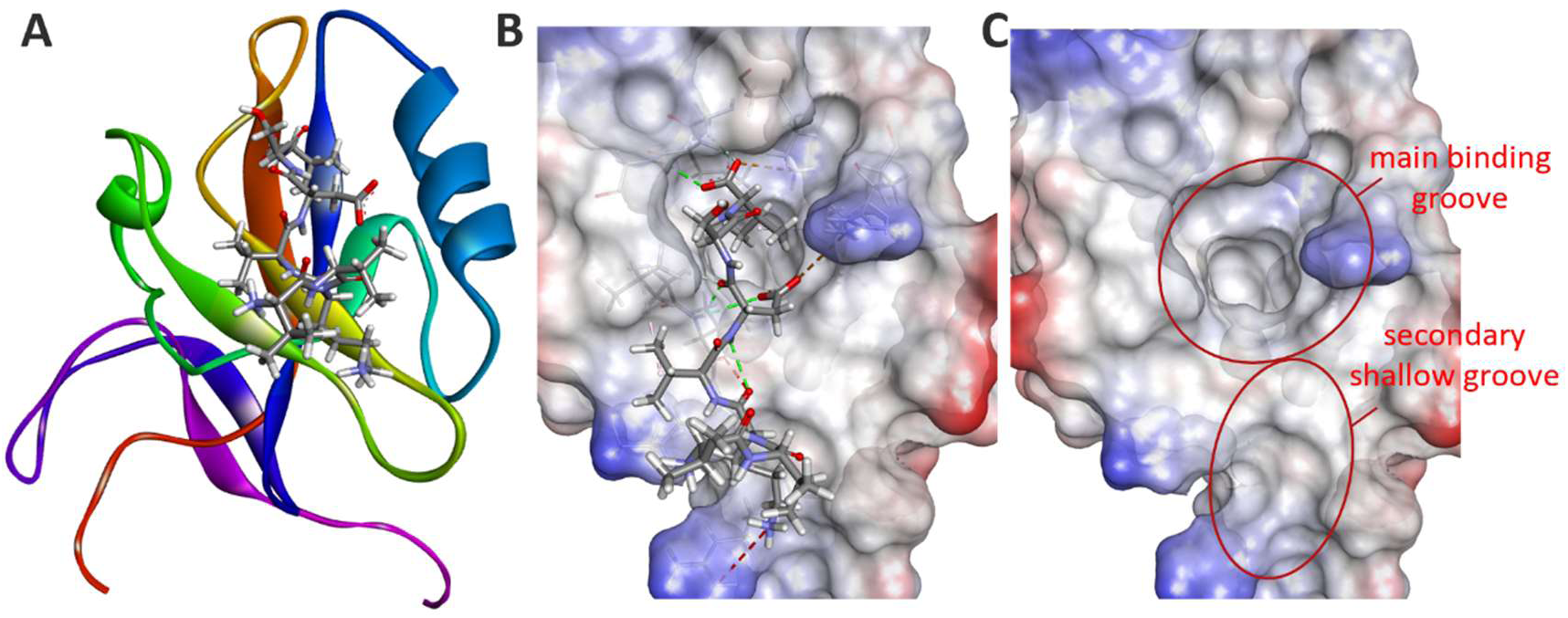
Structural representation of the CAPON-CT7 peptide at the nNOS binding site. (A) Ribbon representation of nNOS showing the CAPON-CT7 peptide (sticks) bound within the protein binding site. (B) Surface representation of the nNOS active site, highlighting the CAPON-CT7 peptide (sticks) and key interactions (dashed lines). (C) Electrostatic surface view of the binding site showing the main binding groove and a secondary shallow groove that may contribute to ligand specificity and affinity.

For a compound to advance as a viable hit candidate, it was required to meet specific structural criteria that recapitulate the binding characteristics of the native CAPON-CT7 interaction. Specifically, each potential hit had to fully occupy the main binding groove of the nNOS/CAPON-CT7 interface, which represents the primary interaction site. In addition, each potential hit was required to demonstrate either partial or complete occupation of the secondary shallow groove (Figure 4C). These stringent geometric requirements ensured that selected compounds would maintain the essential molecular contacts necessary for effective nNOS inhibition. Following this selection process, six compounds met all established criteria and were designated as high-priority hit candidates for further validation.

Analysis of the 2D structures (Figure 5) of the top 6 predicted hits showed a highly consistent pharmacophore profile where all six compounds exhibited peptidomimetic or peptide-like structures which is characterized by containing multiple amide bonds which is essential for protein-protein interaction mimicry. Compounds **1**, **3**, **4**, **5**, and **6** all incorporate aromatic ring systems, predominantly benzene rings. The presence of heterocyclic moieties is another recurring feature, with compounds **2**, **3**, **5**, and **6** containing various nitrogen-containing rings such as pyrrolidines, piperidines, or other cyclic structures. These heterocycles likely contribute to conformational rigidity and may participate in specific hydrogen bonding networks or electrostatic interactions with the protein.

**Figure 5.**
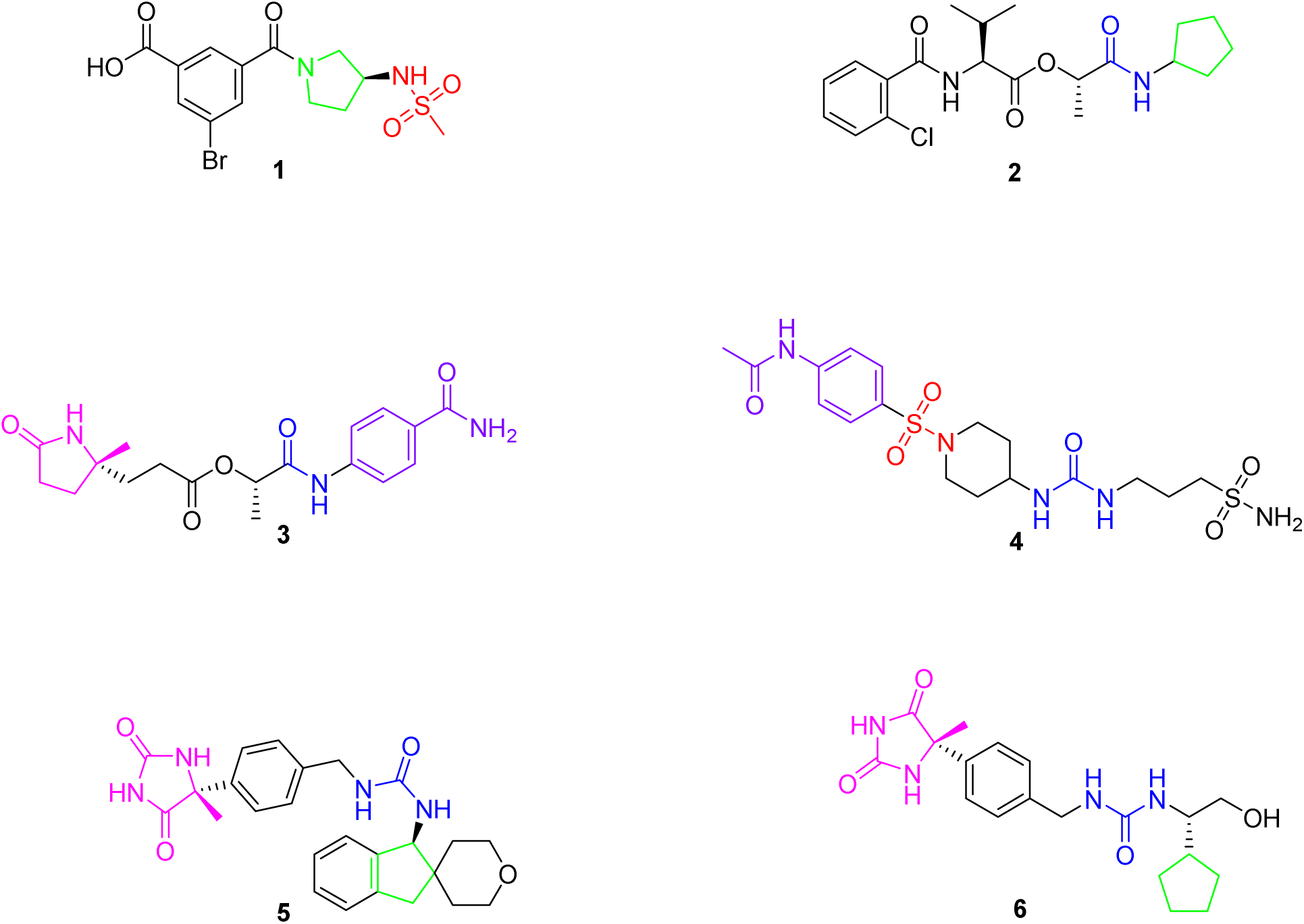
2D structures of the top 6 predicted hits.

Structure flexibility is another common feature which appears to be modulated through the incorporation of linear alkyl chains and ester linkages, as observed in compounds **2**, **4**, and **6**. The varying chain lengths and branching patterns suggest that there is some tolerance for structural diversity in these regions, while maintaining essential pharmacophoric elements. Compounds **1** and **4** contain halogen substituents (bromine and chlorine, respectively as well as sulfur-containing functional groups.

While the virtual screening assay predicted high binding activity and favorable interactions for the hit compounds, this approach is inherently limited by treating the protein as a rigid structure^22^ while allowing only ligand flexibility. To address these limitations and investigate both binding pose stability and protein conformational dynamics under physiologically relevant conditions, molecular dynamics (MD)^23, 24^ simulations were carried out. A total of 8 independent 100 ns MD simulations (Figure 6) were conducted to account for the seven protein-NOS complexes representing each hit compound bound to NOS, the NOS/CAPON-CT7 complex containing the C-terminal 7-residue peptide of CAPON to validate the natural protein-protein interaction and the unbound NOS protein to establish baseline dynamics. This systematic approach enabled assessment of binding pose stability, conformational changes upon ligand binding, and comparison with the native CAPON/NOS interaction dynamics.

**Figure 6.**
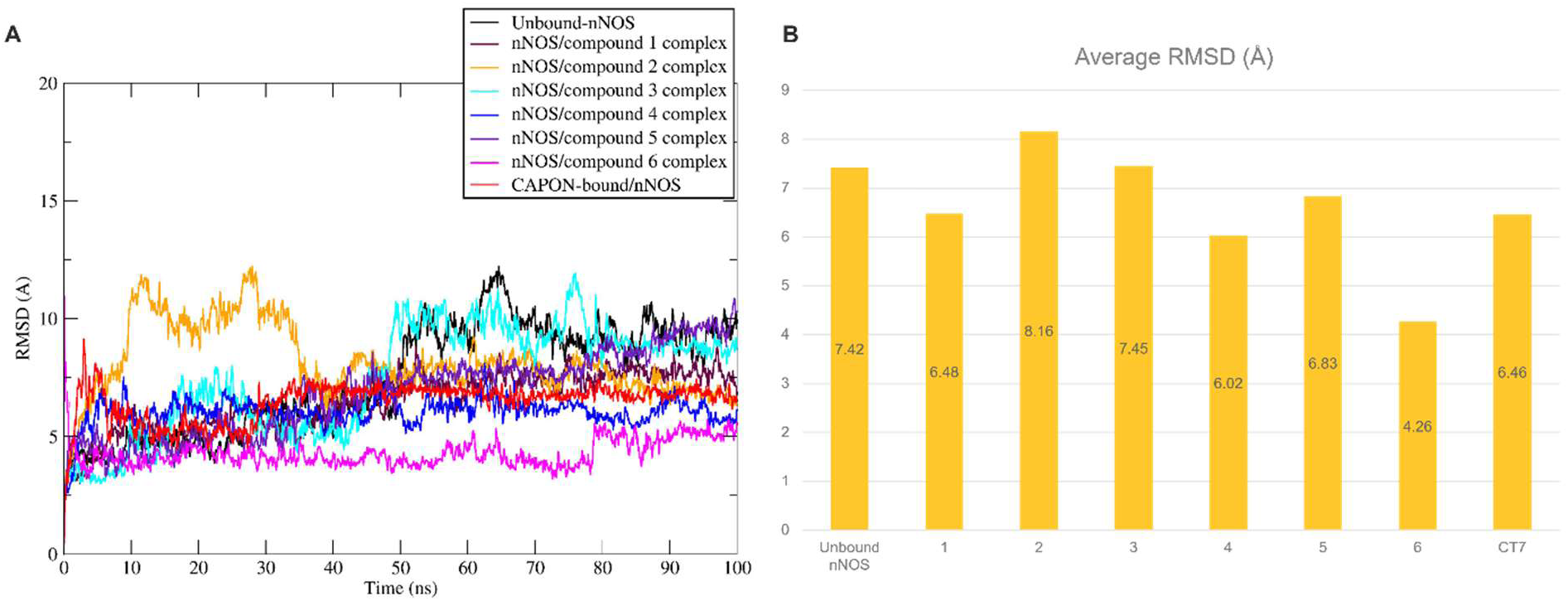
RMSD trajectories during 100 ns molecular dynamics simulations of nNOS protein complexes in comparison with the unbound nNOS protein. (A) RMSD trajectories of nNOS protein backbone atoms in complex with hit compounds **1**-**6**, nNOS/CT7 complex and the unbound nNOS. (B) Comparative plot of the Average RMSD of the established complexes with nNOS.

The stability of the ligand–target complexes was evaluated through root-mean-square deviation (RMSD) analysis^25^. RMSD values (Figure 6) were calculated for both the protein backbone and the bound ligands over the course of the simulation to monitor conformational fluctuations and assess whether the complexes remained stable throughout the trajectory. Consistently low RMSD values indicate minimal structural deviation, thereby supporting the structural integrity of the binding poses and the overall validity of the simulation setup^26^.

Consequently, the RMSD trajectory of the ligand-protein complexes in comparison to the unbound nNOS was carried out. The unbound nNOS protein (Figure 6A, black line) demonstrates relatively high structural fluctuations with an average RMSD of 7.42Å throughout the simulation, indicating significant conformational flexibility in the absence of a binding partner. In contrast, the CAPON-bound nNOS complex (red line) exhibited a more stable profile with average RMSD of 6.46Å, establishing this as the reference state for optimal binding site stabilization.

Among the six predicted hit compounds, compounds **4** (blue) and **6** (pink) demonstrate the most favorable stability profiles by maintaining RMSD values lower than the native CAPON-CT7 complex, with average RMSD values of 6.02Å and 4.26Å, respectively. This enhanced stability could translate to improved binding affinity and longer residence times at the target site. Compounds **1**, **2**, **3**, and **5** showed high fluctuations during the MD simulation, especially compound 3 which exhibited increased fluctuation around the 60-70 ns timeframe. These results suggest that all six compounds possess the capacity to bind with the nNOS binding site, with compounds 4 and 6 emerging as the most promising candidates based on their ability to maintain protein structural integrity during the simulation.

Based on these results, the docked poses of compounds **4** and **6** were chosen to be illustrated in Figure 7 in comparison with the CAPON-CT7 complex. The comparative binding mode analysis revealed distinct interaction patterns between the natural CAPON-CT7 peptide and the three most promising hit compounds. The native NOS/CAPON-CT7 complex (Figure 7A-B) establishes the reference binding profile with hydrogen bonds between CT7 peptide and Leu22, Gly23, Phe24, Val26 and Tyr71 amino acid residues of the Nos binding site. The CT7 peptide adopted an extended conformation.

**Figure 7.**
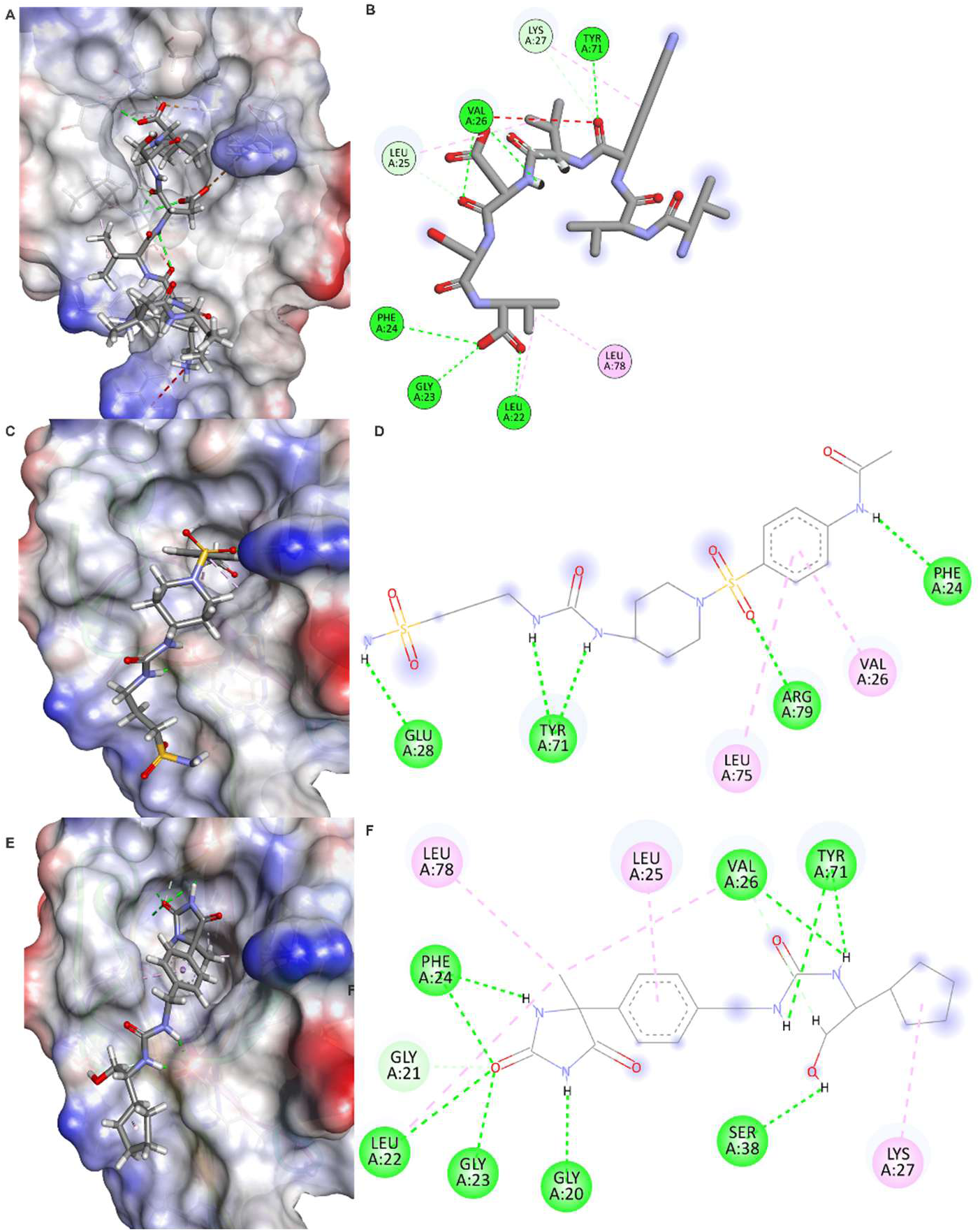
Molecular binding modes and interaction profiles of CAPON-CT7 peptide and hit compounds with NOS protein. (A-B) NOS/CAPON-CT7 natural complex showing the binding pose and interaction network of the C-terminal 7-residue CAPON peptide within the NOS binding pocket. (C-D) NOS/Compound **4** complex displaying the small molecule binding mode and key residue interactions. (E-F) NOS/Compound **6** complex illustrating the binding conformation and interaction pattern of the most stable compound. Left panels (A, C, E) show the surface representation of the NOS protein (colored by electrostatic potential: red = negative, blue = positive, white = neutral) with bound ligands displayed as stick models within the binding cavity. Right panels (B, D, F) present 2D interaction diagrams showing hydrogen bonds (green dashed lines), hydrophobic contacts (pink circles), and other non-covalent interactions between the ligands and surrounding amino acid residues.

Compound **4** (Figure 7C-D) and compound **6** (Figure7E-F) both occupied the binding site with compound **4** establishing hydrogen bonds with Phe24, Glu28, Arg79 and Tyr71. In addition, the benzene moiety of compound **4** established twoVal26 and Leu75 amino acid residues of the nNOs binding site. Although compound 4 exhibits a relatively high number of predicted interactions with the target, its significant exposure to the solvent-accessible surface likely contributes to the pronounced structural instability observed in the RMSD analysis. This solvent exposure may have contributed to the observed lower stability of the ligand–target complex by weakening key interactions and increasing conformational flexibility during the simulation.

Meanwhile, compound **6** exhibited a broader interaction network when compared to compound **4**. Compound **6** established hydrogen bonds with Gly20, leu22, Gly23, Phe24, Val26, Ser38 and Tyr71 amino acid residues of the nNOS binding site. This extensive contact pattern suggests that compound **6** may achieve superior binding affinity through its ability to simultaneously engage multiple residues within the binding pocket which consistent with its high RMSD stability profile observed in the molecular dynamics simulations.

Next, Root Mean Square Fluctuation (RMSF) calculations were carried out to compare the flexibility of the unbound nNOS and bound protein when complexed with hits **4** and **6** (Figure 8). The RMSF (analysis reveals significant differences in protein flexibility patterns between the unbound nNOS (Figure 8A) and the two compound-bound states (Figure 8B-C). The unbound nNOS exhibits substantial structural flexibility throughout the protein, with particularly pronounced fluctuations in several regions with RMSF values reach up to 14 Å. In contrast, both compound **4** and compound **6** dramatically reduce these fluctuations. Compound **6** demonstrates superior stabilization effects, showing more uniform flexibility reduction across the entire protein structure, with particularly effective dampening of the high-mobility regions observed in the unbound state. This flexibility reduction upon compound binding suggests that both hits successfully engage the protein in a manner that constrains conformational dynamics, with compound **6** providing more comprehensive stabilization that may contribute to its enhanced binding affinity and longer residence time at the target site. The MMGBSA^27^ binding free energy calculations were calculated for the complexes of compounds **4** and **6** with NOS and were calculated to be −18.40 and - 25.63kcal/mol, respectively. These results align with the predicted docking scores and RMSD and RMSF values, supporting compound **6** as the more promising hit of the two.

**Figure 8.**
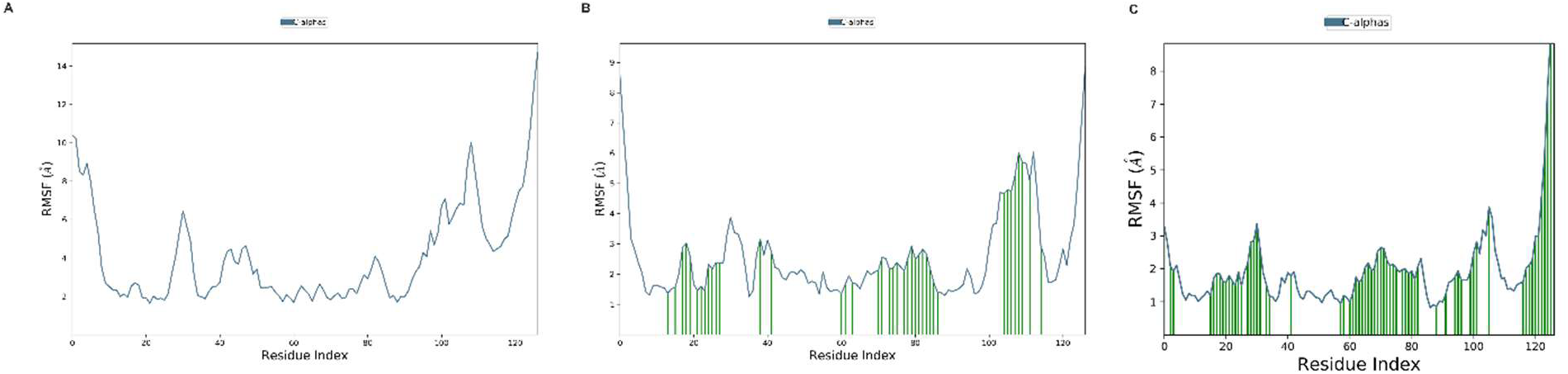
RMSF analysis of nNOS protein flexibility across different binding states. (A) RMSF profile of unbound nNOS showing per-residue atomic fluctuations over the 100 ns molecular dynamics simulation, with high flexibility regions indicating structural mobility in the absence of binding partners. (B) RMSF profile of the nNOS/compound **4** complex demonstrating reduced protein flexibility upon ligand binding, with green bars highlighting regions of significant stabilization compared to the unbound state. (C) RMSF profile of the nNOS/compound **6** complex showing comprehensive protein stabilization with markedly reduced fluctuations across most residues. RMSF values are calculated for Cα atoms and represent the average positional deviation from the mean structure. Lower RMSF values indicate greater structural stability, while higher values suggest increased conformational flexibility. The dramatic reduction in RMSF upon compound binding demonstrates the stabilizing effect of ligand engagement on protein dynamics.

## 3. Methodology

### 3.1. NMR conformations analysis

A python-based script (supplementary file) was developed and employed to analyze the different conformations within the NMR structure of the NOS protein bound to 7 residues of CAPON (PDB ID: 1B9Q^13^). The pipeline processes multi-model PDB files using the MDAnalysis library for structure parsing and manipulation. All conformations are first aligned to a reference structure (the first conformation) using the Kabsch algorithm to minimize positional deviations. The alignment process ensures that subsequent structural comparisons reflect genuine conformational differences rather than rigid-body translations or rotations. Following alignment, a symmetric pairwise RMSD matrix is computed using vectorized operations for all model combinations, where RMSD is calculated as:

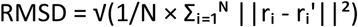

where *N* is the number of atoms, *rᵢ* represents atomic coordinates in the reference structure, and *rᵢ’* represents coordinates in the compared structure.

The script incorporates multiple complementary approaches to characterize conformational diversity. Root Mean Square Fluctuation (RMSF) values quantify per-atom flexibility across the ensemble, while radius of gyration measurements assess the overall structural compactness for each conformation. K-means clustering applied to flattened coordinate vectors identifies distinct conformational states, with cluster numbers adaptively determined based on ensemble size. Dimensionality reduction techniques, including Principal Component Analysis (PCA) for linear variance decomposition and Multidimensional Scaling (MDS) for non-linear distance preservation, provide low-dimensional representations of the conformational landscape. Hierarchical clustering with Ward linkage generates dendrograms that reveal evolutionary relationships between conformations, enabling identification of the most representative (central) structure based on minimum average RMSD to all other models.

Results are exported as structured CSV files containing model statistics and atomic analysis, JSON metadata with summary metrics, aligned PDB structures, and multi-panel visualizations including RMSD heatmaps, statistical distributions, dimensionality reduction plots, and flexibility profiles.

### 3.2. Library preparation

The ligand preparation process was performed using a modular and automated Python pipeline implemented in the chem_pipeline.py script (supplementary file), designed to handle high-throughput molecular preprocessing. Initially, raw ligand structures provided in a delimited text or CSV file were processed using the pipeline’s CSV preparation module. This module automatically detected and standardized the delimiter format (e.g., commas, tabs, or semicolons), ensuring robust compatibility across diverse datasets. Following standardization, the clean CSV was parsed to extract SMILES strings and associated compound IDs, which were then converted into 2D or optionally 3D structure-data file (SDF) representations using RDKit. Error handling and logging were integrated at this stage to discard malformed or non-interpretable SMILES strings.

The resulting SDF was subjected to a comprehensive filtering stage that applied a hierarchy of drug-likeness and safety criteria. Molecules were processed in parallel using multiple CPU cores to accelerate throughput. Filtering included enforcement of Lipinski’s Rule of Five^28, 29^, additional drug-likeness constraints such as topological polar surface area (TPSA), number of rotatable bonds, and fraction of sp³ hybridized carbons, and substructure flags based on PAINS, BRENK, and NIH structural alerts. Canonical tautomers were generated, and molecules were standardized, uncharged, and optionally kekulized and hydrogenated. Molecules failing any criterion were filtered out, and detailed logs were saved for traceability. In the final stage, molecules with non-zero formal charges at physiological PH were identified and removed using a charge screening step, yielding a final SDF file composed exclusively of structurally and physiochemically suitable, neutral ligands. Duplicate structures were eliminated based on canonical SMILES unless explicitly retained by user configuration.

### 3.3. SBVS

Structure-based virtual screening was applied to identify potential CAPON/NOS binding possible modulators from the Enamine compound libraries using a hierarchical docking approach. The NOS protein structure was obtained from the Protein Data Bank (PDB ID: 1B9Q^13^) and prepared for virtual screening using the Protein Preparation Wizard in Schrödinger Suite. The CAPON/NOS binding site was identified and validated through analysis of the co-crystal structure, with receptor grids generated using Glide’s grid generation module to represent the shape and electrostatic features of the binding pocket through multiple separate field sets. The virtual screening protocol employed a three-stage hierarchical filtering approach using Glide’s High-Throughput Virtual Screening (HTVS), Standard Precision (SP), and Extra Precision (XP) docking modes. Initially, the 4.6 million compounds from the Enamine Screening Collection were subjected to HTVS docking. Only the top 10% of compounds from each stage were advanced to the subsequent precision level, ensuring computational efficiency while maintaining screening accuracy. Glide managed conformational flexibility through thorough conformational search followed by rapid exclusion of unfavorable conformations, with ligand poses scored progressively using increasingly accurate algorithms.

### 3.4. Molecular Docking Study

Following the hierarchical screening, compounds that passed the final XP filtering stage were subjected to a molecular docking analysis to predict their binding modes and affinities to the CAPON/NOS binding site. Visual inspection of binding orientations to ensure reasonable protein-ligand interactions and full occupancy was employed to select the most promising hits for further analysis. The selected compounds were re-docked using Glide’s Extra Precision module leading to the generation of multiple poses for each ligand to account for binding mode variability. Subsequently, the docking scores were ranked by selecting the most negative GlideScores for each complex, and binding poses were analyzed for key interactions including hydrogen bonding, hydrophobic contacts, and electrostatic interactions. Molecular visualization and analysis were performed using BIOVIA Discovery Studio Visualizer to assess binding mode feasibility and identify critical pharmacophoric interactions.

### 3.5. Molecular Dynamics study

To validate the stability of predicted binding poses and investigate protein conformational dynamics, molecular dynamics simulations were conducted on the most promising hits. A total of 8 independent 100 ns MD simulations were performed using Desmond molecular dynamics package, including 17 NOS-compound complexes, the unbound NOS protein, and the reference NOS/CAPON-CT7 complex containing the C-terminal 7-residue peptide of CAPON. All systems were solvated in explicit TIP3P water molecules within periodic boundary conditions, with appropriate counter-ions added to maintain system neutrality. The OPLS3e force field was employed for protein and ligand parameterization, with simulations conducted at 300 K and 1 atm pressure using NPT ensemble conditions. Root Mean Square Deviation (RMSD) values were calculated for protein backbone atoms throughout the simulation trajectories to assess binding pose stability and protein conformational changes, with the initial docked structure serving as the reference frame for RMSD calculations. The protein-ligand interactions were analyzed using the Protein Interaction Wizard within DEMSOND. RMSD plots were visualized using the XmGrace software.

### Conclusions

Our virtual screening campaign evaluated 4.6 million compounds to identify novel small molecule modulators of the CAPON/nNOS protein-protein interaction. The virtual screening predicted 6 promising candidates that were subsequently validated using 100 ns molecular dynamics simulations. Comparative analysis against both the unbound protein and the CAPON/NOS complex revealed two lead compounds demonstrating superior binding stability, sustained molecular contacts, and energetically favorable binding conformations. These computationally validated hits represent high-priority candidates for experimental validation, which will commence upon completion of our laboratory’s ongoing efforts to establish a robust biochemical screening platform for CAPON. Beyond the identification of potential therapeutic leads, this work introduces two Python-based toolkits for protein structure analysis and streamlined ligand preparation. The NMR analysis toolkit facilitates rapid processing and visualization of structural data, while the ligand preparation workflow automates and standardizes compound preparation protocols, thereby minimizing manual intervention and potential sources of error. The toolkits significantly reduce computational overhead and processing time compared to conventional approaches which enhances the efficiency of large-scale virtual screening endeavors. Collectively, these findings establish a solid foundation for the rational design of CAPON/nNOS modulators and provide valuable computational tools that will accelerate future drug discovery efforts targeting this therapeutically relevant protein-protein interaction.

## Acknowledgments

We acknowledge funding from the National Institutes on Aging (NIA) RF1AG084635 (PI: Gabr).

## Data Availability Statement

The ligand preparation Python pipeline is provided in the Supplementary Information. The Enamine libraries used for screening are freely accessible through the Enamine server and can be downloaded using the corresponding compound IDs. Additionally, the supplementary folder contains the 4,750 initially predicted hits along with their associated docking scores.

